# Phosphatidylinositol-3 kinase activity controls survival and stemness of zebrafish hematopoietic stem/progenitor cells

**DOI:** 10.1101/2020.07.17.208678

**Authors:** Sasja Blokzijl-Franke, Bas Ponsioen, Stefan Schulte-Merker, Philippe Herbomel, Karima Kissa, Suma Choorapoikayil, Jeroen den Hertog

## Abstract

Hematopoietic Stem and Progenitor Cells (HSPCs) are multipotent cells giving rise to all blood lineages during life. HSPCs emerge from the ventral wall of the dorsal aorta (VDA) during a specific timespan in embryonic development through endothelial hematopoietic transition (EHT). We investigated the ontogeny of HSPCs in mutant zebrafish embryos lacking functional pten, an important tumor suppressor with a central role in cell signaling. Through in vivo live imaging, we discovered that in *pten* mutant embryos a proportion of the HSPCs died upon emergence from the VDA, an effect rescued by inhibition of phosphatidylinositol-3 kinase (PI3K). Surprisingly, inhibition of PI3K in wild type embryos also induced HSPC death. Surviving HSPCs colonized the caudal hematopoietic tissue (CHT) normally and committed to all blood lineages. Single cell RNA sequencing indicated that inhibition of PI3K enhanced survival of multi-potent progenitors, whereas the number of HSPCs with more stem-like properties was reduced. At the end of the definitive wave, loss of Pten caused a shift to more restricted progenitors at the expense of HSPCs. We conclude that PI3K signaling tightly controls HSPCs survival and both up- and downregulation of PI3K signaling reduces stemness of HSPCs.

**Key points:**

- Loss of Pten and inhibition of PI3K induced apoptosis of hematopoietic stem/progenitor cells upon endothelial to hematopoietic transition
- Surviving hematopoietic stem/progenitor cells committed to all blood lineages but displayed reduced stemness

## Introduction

Stem cells define a particular type of cells that maintain self-renewal capacity and may differentiate into multiple cell types at the same time. HSPCs are multipotent cells giving rise to all blood lineages during life^1–3^. In all vertebrates, an initial primitive wave of hematopoiesis occurs in the embryo, giving rise to primitive erythrocytes and myeloid cells. A definitive wave follows in which HSPCs are generated that will found multi-lineage hematopoiesis in developmentally successive hematopoietic organs up to adulthood. Our understanding of the emergence of HSPCs during the definitive wave is derived primarily from pioneer live *in vivo* imaging^4–6^. HSPCs emerge in a process whereby cells in the ventral wall of the dorsal aorta (VDA) undergo an endothelial to hematopoietic transition (EHT)^5^ and then transiently colonize the fetal liver in mammals^7^, or the caudal hematopoietic tissue (CHT) in zebrafish^8^. There, HSPCs expand and differentiate into all blood lineages and supply the developing embryos with mature blood cells. Subsequently, HSPCs migrate again to colonize the thymus and the bone marrow in mammals ^7^ or whole kidney marrow in fish^8^ to produce blood cells in the adult.

HSPCs are tightly regulated in terms of dormancy, self-renewal, proliferation and differentiation. Disrupting this balance can have pathological consequences such as bone marrow failure or hematologic malignancy. The tumor suppressor, PTEN, has an important role in hematologic malignancies, particularly T-lineage acute lymphoblastic leukemia (T-ALL). Deleterious mutations in *PTEN* appear in 5-10% of T-ALL cases and about 17% of patients lack PTEN expression in the hematopoietic lineage^9,10^. PTEN counteracts phosphatidylinositol-3 kinase (PI3K) and hence acts upstream in the PI3K-Akt (also known as Protein kinase B, PKB) pathway^11^. Loss of PTEN function results in hyperactivation of the PI3K-Akt signaling pathway. Clonal evolution of leukemia-propagating cells in zebrafish highlights the role of Akt signaling in the process^12^. Conditional deletion of Pten in mice in hematopoietic stem cells (HSCs) in adult bone marrow promotes HSC proliferation, resulting in depletion of long-term HSCs, indicating that Pten is essential for the maintenance of HSCs^13,14^.

The zebrafish genome encodes two *pten* genes with redundant function designated *ptena* and *ptenb*^15^. Single mutants display no morphological phenotype and are viable and fertile, but mutants that retain only a single wild type copy of *pten* develop hemangiosarcomas during adulthood^16^. *Ptena^−/−^ptenb^−/−^* mutants lack functional Ptena and Ptenb, are embryonic lethal at 5-6 days post fertilization (dpf) and display hyperplasia and dysplasia^15^. We reported that double mutant zebrafish larvae lacking functional Pten display increased numbers of HSPCs in the CHT at 4-5 dpf. Whereas these HSPCs commit to different blood lineages, they fail to differentiate into mature blood cells. Inhibition of PI3K using LY294002, which compensates for the loss of Pten, restores differentiation of HSPCs into mature blood cells. Hence loss of Pten enhances HSPCs proliferation and arrests differentiation^17^.

The past decades have led to an increase in our knowledge of hematopoiesis, but we are still far from a complete understanding of how HSPCs are established. Likewise, the role of Pten in steady-state hematopoiesis has been studied, but its potential role in the ontogeny of HSPCs is not fully understood. We have addressed these questions in zebrafish larvae. We imaged the emergence of HSPCs from the VDA *in vivo* in *ptena^−/−^ptenb^−/−^* embryos and in PI3K-inhibitor treated wild type embryos, which showed surprisingly similar defects. Furthermore, we performed single cell RNA sequencing (scRNA-seq) during the onset and at the end of the definitive wave. Our results indicate that elevated and reduced PI3K signaling had opposite effects on HSPCs at the end of the definitive wave.

## Methods

### Ethics statement

All animal experiments described in this manuscript were approved by the local animal experiments committee (Hubrecht Institute: Koninklijke Nederlandse Akademie van Wetenschappen-Dierexperimenten commissie protocol HI180701 and University Montpellier: Direction Sanitaire et Vétérinaire de l’Hérault and Comité d’Ethique pour l’Expérimentation Animale under reference CEEA-LR-13007) and performed according to local guidelines and policies in compliance with national and European law.

### Zebrafish husbandry

*Ptena^−/−^ptenb^−/−^, ptena^−/−^, ptenb^−/−^*^15^ *, Tg(kdrl:eGFP)*^18^ *, Tg(kdrl:mCherry-CAAX)*^19^, and *Tg(cd41:eGFP)*^20^ were maintained according to FELASA guidelines, crossed, raised and staged as described^21–23^. *Pten* mutant fish (embryos) were genotyped by sequencing^15^. The *tg(kdrl:Dendra2)* line was derived by Tol2-mediated transgenesis^24^ of a construct containing the ~7.0kb *kdrl-*promoter (a kind gift from D. Stainier), driving the expression of Dendra2^25^. From 24hpf onwards, all embryos were grown in PTU-containing medium to block pigmentation

### LY294002 treatment

Embryos were treated with 5 μM LY294002 (Calbiochem, San Diego, CA, USA) or DMSO control in the dark. For early treatment, embryos were incubated with LY294002 from 32 hpf onwards and mounted after 4 hours for time-lapse confocal imaging. For late treatment and to investigate thymus and kidney colonization, embryos were treated with 5 μM LY294002 from 32 to 60 hpf and imaged.

### Confocal, fluorescence, brightfield microscopy and time-lapse imaging

Fluorescence images of transgenic embryos were acquired using TCS-SPE and time-lapse imaging using TCS-SP2 as described^26^ and processed with ImageJ^27^. For all live imaging embryos were anesthetized with tricaine^21^, mounted on a glass cover dish with 0.7% low melting agarose and covered with standard E3 medium. Whole mount bright field images were taken with a Leica DC 300F stereomicroscope.

### *In situ* hybridization

Whole mount *in situ* hybridization was performed according to standard protocols^28^ and images were taken using a Zeiss Axioplan microscope connected to a Leica DFC480 camera.

### Acridine orange staining and whole mount immunohistochemistry

Embryos were incubated with 5 μg/ml acridine orange^29^ for 20 minutes between 35 and 40 hpf and subsequently washed with standard E3 medium. Embryos were then imaged as described above. Immunohistochemical labeling performed using fixed (40 hpf) embryos to detect apoptosis using an activated caspase-3-specific antibody (BD Pharmingen)^30^. After confocal images were collected embryos were genotyped.

### Photoconversion

Fluorescent tracing of VDA-derived HSPCs colonizing the CHT was done using the *tg(kdrl:Dendra2)* line as described before^31,32^ with a Leica SP5 confocal microscope with a 20x dry objective. At 28 hpf an area of approximately 40×750 nm around the VDA, parallel to the yolk sac extension was photoconverted. The 405nm UV laser intensity and exposure time were optimized for strong Dendra2-conversion without cell damage. After photoconversion embryos were transferred to E3 medium and at 50-60 hpf their CHT areas were imaged on a Leica SPE Live confocal microscope using a 20x dry objective. To exclude bleed-through of Dendra-green, red channel detection was set stringently (630-680nm).

### Quantification of GFP^low^ progenitor cells using *tg(cd41:eGFP)*

GFP^low^ and GFP^high^ expressing cells were quantified in the CHT at 48 hpf, 50 hpf or 4 dpf using confocal imaging and Volocity and Imaris software. *Ptena^+/−^ptenb^−/−^* mutants on *tg(cd41:eGFP)* background were crossed and offspring was mounted at 48 hpf. Wild type *tg(cd41:eGFP)* embryos were treated with 5 μM LY294002 as described above and mounted and imaged at 50 hpf or 4 dpf. All GFP^low^ expressing cells were counted in the entire CHT.

### Flow cytometry

The AGMs of approximately 4000 36 hpf and 400 CHTs of 5 dpf old *tg(kdrl:mCherry/cd41:eGFP)* embryos were dissected and collected in Leibovitz-medium. After washing with PBSO the AGMs were deyolked using calcium-free Ringer’s solution (116mM NaCl, 2.9mM KCl and 5mM HEPES) and then AGMs and CHTs were dissociated in TrypLE Express (Gibco) for 45 minutes at 32°C. The resulting cell suspension was washed in PBSO and passed through a 40-μm filter after resuspension in PBS supplemented with 2mM EDTA, 2% FCS and 0.5μg/ml DAPI. DAPI staining was used to exclude dead cells. Cells with kdrl- and cd41^low^-positive signal were subjected to fluorescence-activated cell sorting (FACS) using a BD FACSAriaII and BD FACSFusion.

### ScRNA-seq with SORT-seq

ScRNA-seq was performed by Single Cell Discoveries BV (Utrecht, the Netherlands), according to an adapted version of the SORT-seq protocol^33,34^, with adapted primers described in^35^.

### Data analysis

During sequencing, Read1 used for identification of the Ilumina library barcode, cell barcode and UMI. Read2 was used to map to the reference transcriptome of Zv9 *Danio rerio*. Data was demultiplexed as described^36^. Single cell transcriptomics analysis was done using the RaceID3 algorithm, following an adapted version of the RaceID manual (https://cran.r-project.org/web/packages/RaceID/vignettes/RaceID.html). Cells that had less than 1500 UMIs and genes that were detected in less than 5 UMIs in 5 cells were discarded. The number of initial clusters was set at 5. Differential gene expression analysis was done as described in^33^ with an adapted version of the DESseq2 algorithm^37^.

### Data sharing statement

For original data, please contact.

scRNA data are avaliable at GEO under accession number XXXX

## Results

### The onset of the definitive wave of hematopoiesis is independent of Pten

The onset of the definitive wave starts at 32 hours post fertilization (hpf) with the specification of endothelial cells that will become HSPCs in the floor of the dorsal aorta (DA) in the aorta-gonad-mesonephros (AGM) region (Figure 1a), a conserved process between mammals and zebrafish^4–6^. *Runx1* expression from 32 hpf onwards and *c-myb* expression from 35 hpf onwards mark the hemogenic endothelium of the VDA and its HSPC progeny^8,38^. We found that *ptena^−/−^ptenb^−/−^* mutant embryos expressed *runx1* and *c-myb* along the VDA during the period that HSPCs emerge (between 30 and 44 hpf) just like their siblings (Figure S1), indicating that loss of Pten does not affect initiation of definitive hematopoiesis.

**Fig. 1.**
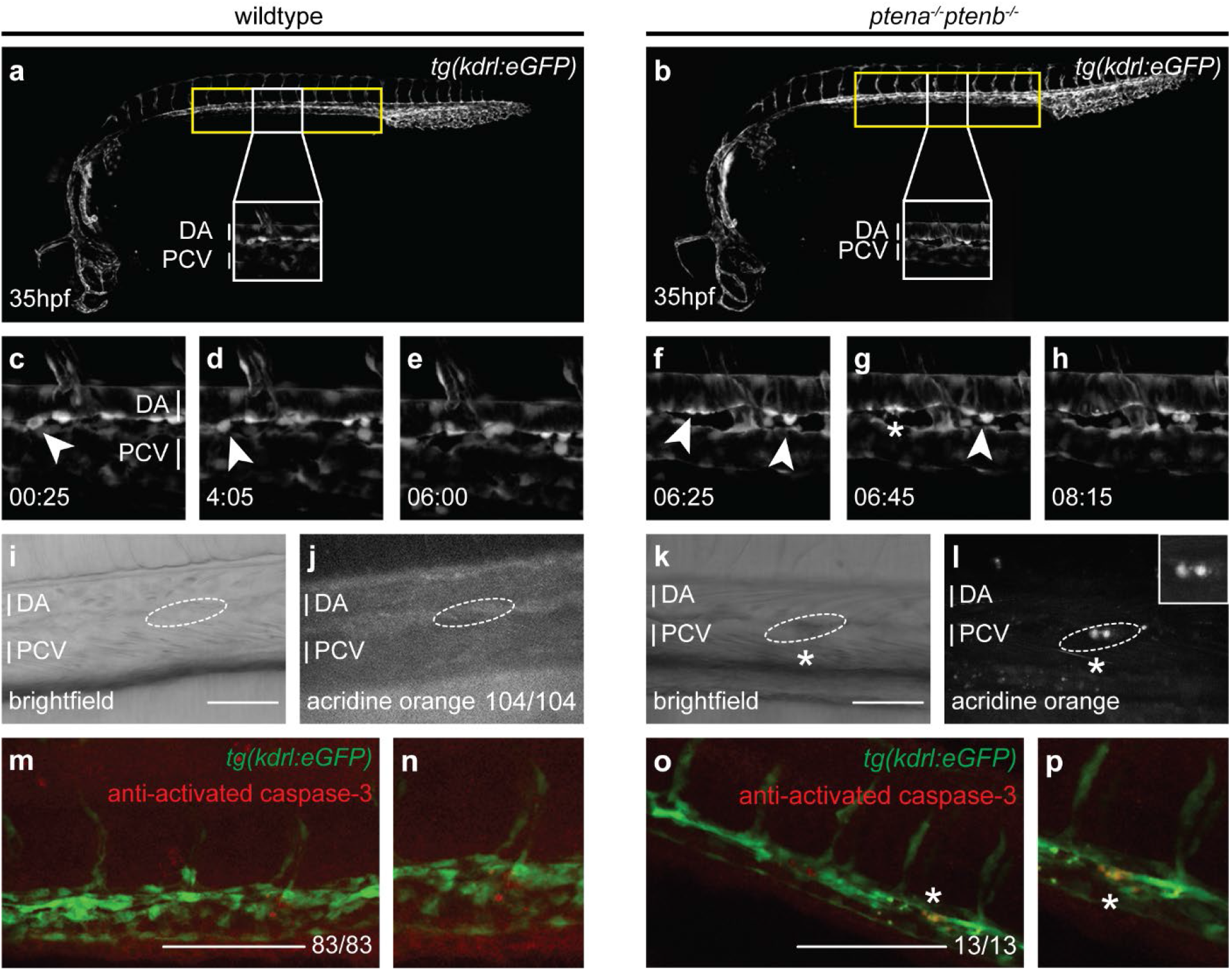
A population of HSPCs fails to complete EHT and undergoes apoptosis in *ptena^−/−^ptenb^−/−^* mutant embryos. (a, b) Brightfield image of a wild type or *ptena^−/−^ptenb^−/−^* mutant zebrafish embryo at 35 hpf. The area from which HSPCs originate is indicated with a yellow box. A close up is indicated with a white box. (c-h) Four-dimensional imaging of *tg(kdrl:eGFP)* wild type or *ptena^−/−^ptenb^−/−^* mutant embryos between 35 hpf and 48 hpf. Arrowheads: HSPCs undergoing EHT; asterisk: disintegrating HSPCs. Confocal image z-stacks (2μm step size, with 40x objective and 2x zoom; anterior to the left; maximum projections of a representative embryo; time in hh: mm. (I-l) Acridine orange staining. Arrows and circles: HSPCs in VDA of 40-45 hpf embryos. Asterisks: apoptotic HSPCs. Scale bar: 50μm. Representative embryos are shown and the number of embryos that showed this pattern/total number of embryos is indicated. DA: dorsal aorta; PCV: posterior cardinal vein. (m-p) confocal images of apoptotic endothelial cells in the VDA of fixated wild type or *ptena^−/−^ptenb^−/−^* mutant zebrafish embryos. In green: *tg(kdrl:eGFP)*; in red: anti-activated caspase-3 immunohistochemistry staining. Apoptotic cells are indicated with an asterisk. Representative embryos are shown and the number of embryos displaying this particular pattern/total number of embryos is indicated in the bottom right. Anterior to the left; 2μm step size; maximum projections; scale bar: 100μm.

### Loss of Pten results in apoptosis of HSPCs during EHT in *ptena^−/−^ptenb^−/−^* mutant embryos

In zebrafish, endothelial cells from the VDA transform into HSPCs in a process called EHT^5^. Subsequently, HSPCs join the blood flow in the underlying posterior cardinal vein (PCV) to transiently seed the CHT^4,5,8^. We monitored EHT events in the AGM by time-lapse confocal imaging of an area spanning two adjacent intersegmental vessels in the tg*(kdrl:eGFP)* transgenic background from 35 to 48 hpf (Figure 1a,b). The vasculature of *ptena^−/−^ptenb^−/−^* mutants and siblings was indistinguishable at this stage^39^. The floor of the aorta in *ptena^−/−^ptenb^−/−^* mutant embryos displayed the characteristic contraction then bending of cells towards the subaortic space^5^, indicating that the initiation of EHT was normal in *ptena^−/−^ptenb^−/−^* mutant embryos. However, half of the EHT events in *ptena^−/−^ptenb^−/−^* mutant embryos were abortive, in that 13 out of 24 HSPCs (54% in 9 embryos) failed to detach and disintegrated (Figure 1c-h, Movie S1). In contrast, siblings or wild type embryos did not display abortive EHT (n=75 in total). Live imaging using acridine orange^29^ revealed apoptotic cells in the DA of *ptena^−/−^ptenb^−/−^* mutant embryos, but not siblings (Figure 1i-l). Activated-caspase-3 immunostaining^30^ confirmed apoptosis of *kdrl:eGFP*-positive cells at the VDA in *ptena^−/−^ptenb^−/−^* mutant embryos (Figure 1m-p). Hence, about half of the HSPCs in *ptena^−/−^ ptenb^−/−^* mutant embryos failed to complete EHT and instead underwent apoptosis.

### The number of HSPCs that colonize the CHT is reduced in *ptena^−/−^ptenb^−/−^* mutant embryos

Following EHT, HSPCs transiently colonize the CHT^8^. We generated a tg*(kdrl:Dendra2)* transgenic line. The Dendra2 protein along the entire VDA was photoconverted green-to-red between 26 and 28 hpf, *i.e.* before the onset of EHT events (Figure 2a). Photoconverted HSPCs in *ptena^−/−^ptenb^−/−^* mutant embryos colonized the CHT between 50 and 60 hpf, albeit less HSPCs were detected than in the CHT of siblings (Figure 2b,c). We quantified the number of HSPCs that colonized the CHT at 48 hpf, *i.e.* by the peak of HSPC emergence from the VDA, using tg*(cd41:eGFP)* embryos, which express low GFP (GFP^low^) in HSPCs^20,40^. Consistent with the initial apoptosis of half of the EHT derived HSPCs, 51% less GFP^low^ HSPCs were detected in the CHT of *ptena^−/−^ptenb^−/−^* mutant embryos at 48 hpf compared to siblings (Figure 2d-f, Figure S2).

**Fig. 2.**
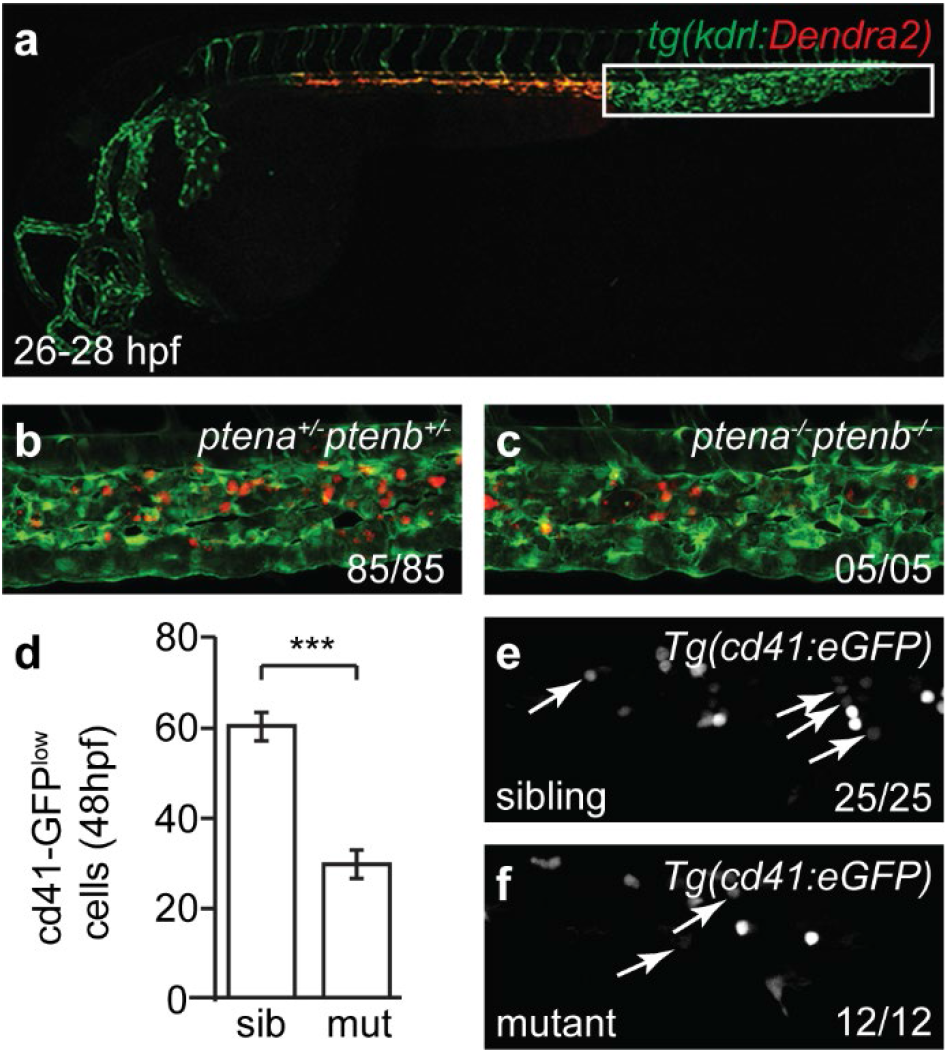
Less HSPCs colonize the CHT in *ptena^−/−^ptenb^−/−^* mutant embryos than in wild type. (a) The VDA of *tg(kdrl:Dendra2)* was photoconverted green-to-red at 26-28 hpf. By 50-60 hpf red HSPCs derived from the photoconverted VDA had colonized the CHT in (b) sibling and (c) *ptena^−/−^ptenb^−/−^* larvae. (d) The number of GFP^low^ HSPCs at 48 hpf in the CHT of *tg(cd41:eGFP)* siblings (sib)(n=25) (e) and *ptena^−/−^ptenb^−/−^* mutants (mut) (n=12) (f) is expressed as average number of cells. Error bars indicate standard error or the mean (SEM). Shapiro Wilk Test for normal distribution and two-tailed t-test were used for statistical analysis; ***p<0.001. Representative embryos are shown and the number of embryos that showed this pattern/total number of embryos is indicated.

### PI3K inhibition rescues EHT events in *ptena^−/−^ptenb^−/−^* mutant embryos but is detrimental for HSPCs in wild type embryos

To address whether apoptosis of half of the EHT-derived HSPCs was due to enhanced PI3K signaling, embryos were treated with the PI3K inhibitor LY294002 from the onset of EHT (32 hpf) onwards. Inhibition of PI3K prevented apoptosis of HSPCs in *ptena^−/−^ptenb^−/−^* mutant embryos, in that none of the HSPCs that we imaged disintegrated (table 1) (Fisher’s exact test, p=0.0013) (Figure 3a-c). Surprisingly, in wild type and sibling embryos that were treated with LY294002 in parallel with the *ptena^−/−^ptenb^−/−^* mutant embryos, disintegrating HSPCs in the VDA were observed (n=6) (Figure 3d-g, Movie S2) (Fisher’s exact test, p=0.021).

**Table 1.** Number of apoptotic cells during EHT. The number of cells undergoing apoptosis during EHT was determined in control and LY294002 (5μM from 32 hpf onwards) treated wild type and *ptena^−/−^ptenb^−/−^* embryos (35-48 hpf) by confocal time lapse imaging. The number of embryos that was imaged is given as well as the number of apoptotic HSPCs or surviving HSPCs. The percentages of emerging or apoptotic HSPCs, relative to the total number of HSPCs are also indicated.

|  |  |  | HSPCs |  |  |  |
| --- | --- | --- | --- | --- | --- | --- |
|  |  | embryos (n) | apoptotic | surviving | % apoptotic | % surviving |
| Wild type | control | 13 | 0 | 24 | 0 | 100 |
|  | LY294002 | 6 | 6 | 9 | 40 | 60 |
| *ptena*<sup>-/-</sup><br>*ptenb*<sup>-/-</sup> | control | 9 | 13 | 11 | 54 | 46 |
|  | LY 294002 | 5 | 0 | 21 | 0 | 100 |

**Fig. 3.**
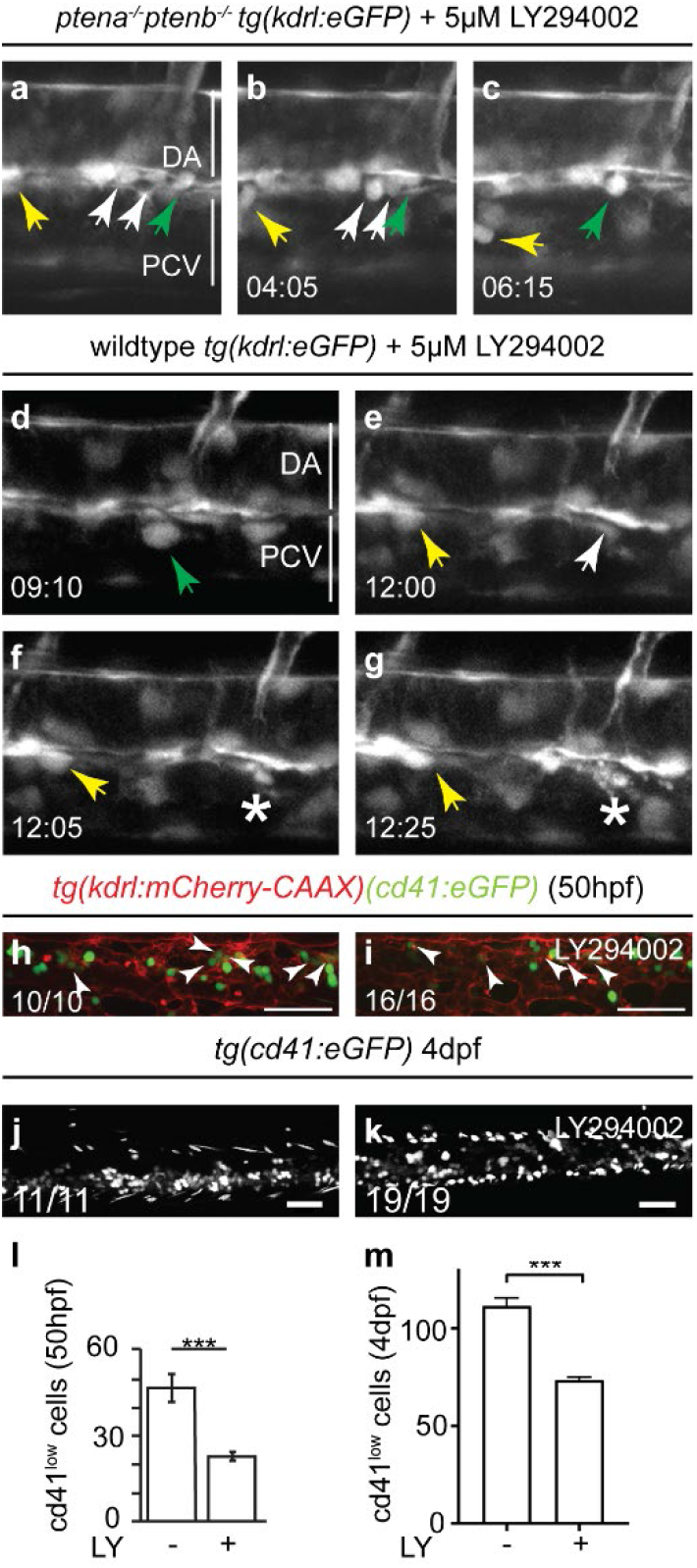
Inhibition of PI3K rescued EHT in *ptena^−/−^ptenb^−/−^* mutant embryos, but induced abortive EHT in wild type embryos. (a-g) Four-dimensional imaging of *tg(kdrl:eGFP)* transgenic embryos. (a-c) *ptena^−/−^ ptenb^−/−^* mutant embryos and (d-g) wild type embryos. Imaging was done from 35 hpf onwards following treatment with 5 μM LY294002 from 32 hpf. Arrowheads: HSPCs. Asterisks: disintegrating HSPCs. Different colors of arrowheads distinguish separate EHT events. Images were taken with 40x objective and 1x zoom. Time in hh:mm; DA: dorsal aorta; PCV: posterior cardinal vein. (h, i) CHTs of *tg(kdrl:mCherry-CAAX/cd41:eGFP)* control (n=10) and LY294002 treated (5 μM, 32-50 hpf) (n=16) embryos were imaged at 50 hpf. The vasculature is highlighted in red (mCherry) and some GFPlow HSPCs are indicated by arrows. (j-k) CHTs of tg(cd41:eGFP) control(n=11) and LY294002-treated (5μM, 30-60 hpf) (n=19) embryos were imaged at 4 dpf. Anterior to the left; 2 μm step size. Representative embryos are shown and the number of embryos that showed this pattern/total number of embryos is indicated. (l-m) the number of GFP^low^ HSPCs was determined at 50 hpf (l) and 4 dpf (m) and is expressed as average number of cells; error bars indicate standard error of the mean (SEM). Shapiro Wilk Test for normal distribution and two-tailed t-test were used for statistical analysis; *** p<0.001.

Consistent with abortive EHT events upon LY294002 treatment, significantly less GFP^low^ HSPCs in *tg(cd41:eGFP)* transgenic embryos colonized the CHT of LY294002-treated wild type embryos at 50 hpf (Figure 3h-i,l), comparable to the reduction observed in *ptena−/−ptenb−/−* mutant embryos (Figure 2d). This reduction of HSPCs persisted through 4 dpf in LY294002-treated embryos (Figure 3j-k,m). These data suggest that normal, i.e. not too high and not too low PI3K activity levels are essential for emergence of HSPCs.

### PI3K inhibition in wild type embryos results in HSPCs that engage in all blood lineages

*In situ* hybridization was performed using a panel of blood progenitor markers. LY294002-treatment reduced expression of the HSPC marker *c-myb* at 4 dpf (Figure 4a,b). The lineage markers *globin* (erythroid lineage), *ikaros* (lymphoid lineage) and *l-plastin* (pan-leukocytic, including myeloid lineage) were expressed, but reduced in LY294002-treated larvae compared to controls (Figure 4c-h). LY294002-treated HSPCs also committed to the thrombocytic lineage as demonstrated by GFP^high^ cells in tg*(cd41:eGFP)* embryos at 5 dpf^20^ (Figure 4i,j). The number of GFP^low^ cells in the definitive hematopoietic organs, thymus and kidney of at 8 and 12 dpf *tg(cd41:eGFP)* larvae was reduced in response to LY294002 (Figure 4k-n). These results show that reduced PI3K signaling did not block specification of particular blood lineages, but that the reduction in HSPC numbers affected founding of the definitive hematopoietic organs by HSPCs.

**Fig. 4.**
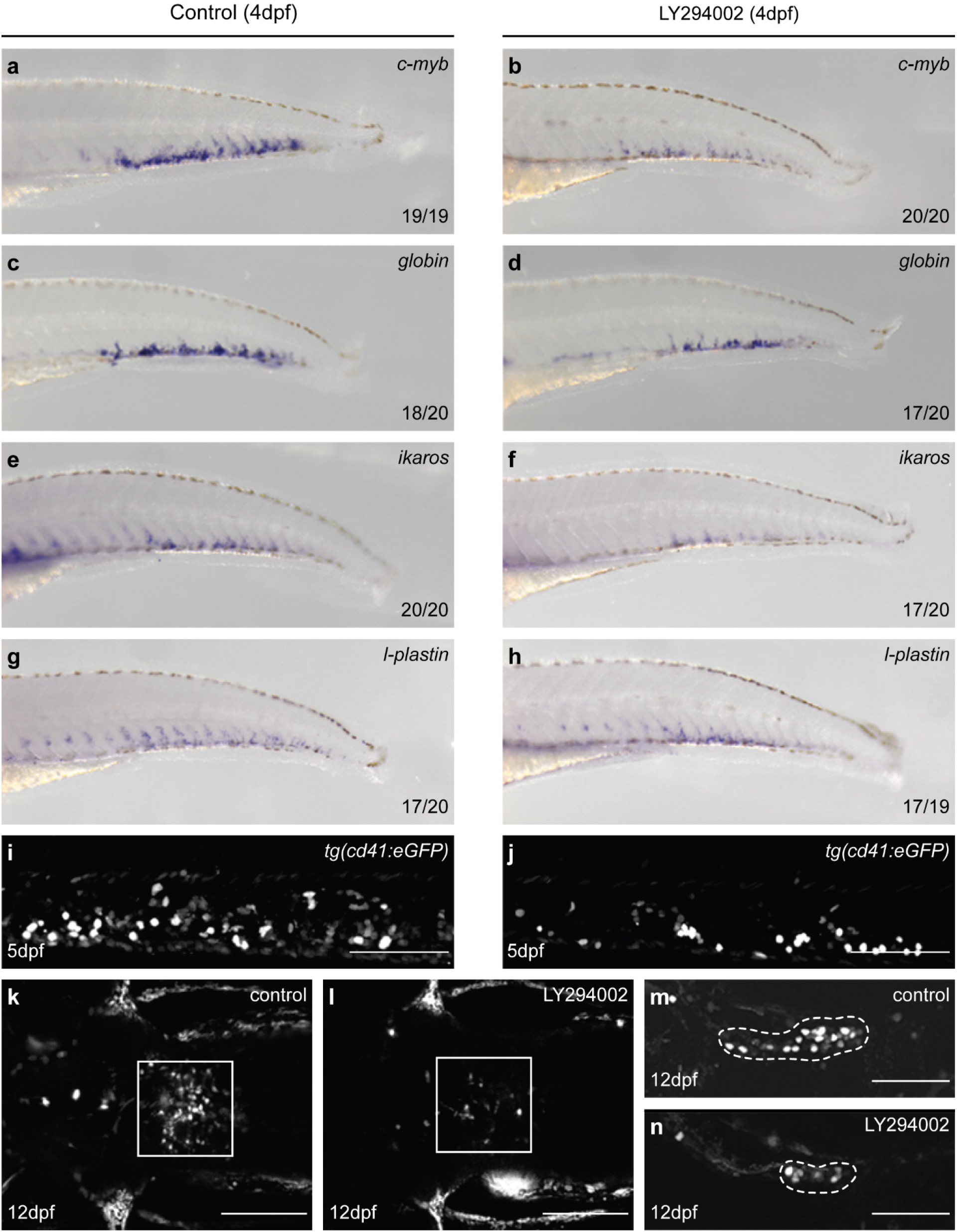
HSPCs of LY294002-treated embryos engage in all blood lineages, but show impaired colonization of definitive hematopoietic organs. (a-h) Control and LY294002-treated (from 32-60 hpf) embryos were fixed at 4 dpf. Markers for definitive blood lineages were assessed by *in situ* hybridization in the CHT: *c-myb* (HSPCs; a,b), *globin* (erythrocyte lineage; c,d), *ikaros* (lymphocyte lineage; e,f), *l-plastin* (leukocytes; g,h). Representative embryos are shown, with anterior to the left. The number of embryos that showed a particular pattern/total number of embryos is indicated in the bottom right corner of each panel. (i-j) GFP^high^ thrombocytes were imaged at 5dpf in tg(cd41:eGFP) embryos. Scale bar: 100μm. (k-n) High-resolution imaging at 12 dpf of kidney (l, m, dorsal view) (control, n=6; LY294002-treated, n=8; 4 μm step size) and thymus (n, o, lateral view) (control, n=6; LY294002-treated, n=7; 2 μm step size). Anterior to the left; maximum projections of representative larvae. Scale bar: 100 μm.

### Singe cell RNA sequencing reveals two types of HSPCs, one of which is preferentially lost upon inhibition of the PI3K-pathway

To investigate transcriptomic changes in HSPCs between LY294002-treated embryos and their controls during EHT, we performed scRNA-seq. Transgenic tg*(kdrl:mCherry-CAAX/cd41:eGFP)* embryos were treated with LY294002 and AGM regions were isolated by dissection at 36hpf. The AGM regions of approximately 2000 control embryos were pooled and likewise, 2000 LY294002-treated embryos. The cells were dissociated and sorted for mCherry^+^/ eGFP^low^ using FACS, after which the SORT-Seq protocol was performed^33^(Figure S3). Isolation of 3,219 cells in total, *i.e*. less than one kdrl^+^/cd41^low^ (mCherry^+^/GFP^low^) HSPC per embryo, was in line with earlier reports (3 HSPCs per embryo per hour^5,41^). After FACS filtering, 2512 cells remained. RaceID3^42^ was used for differential gene expression analysis and clustering of the cells (Figure 5). The resulting t-SNE map highlighted particular cell types, in line with recent scRNA-seq studies of hematopoietic organs of zebrafish^43–50^, which expressed validated hematopoietic lineage markers (Table S1). Cells in cluster 2 and 4 expressed HSPC-related genes, such as *gata2b, gfi1aa, meis1b, myb* and *pmp22b,* consistent with expression in mammalian HSCs and zebrafish HSPCs^44–48,50^ (Figure 5a,b). RaceID3 subdivided the main HSPC cluster into two, HSPCs I and HSPCs II, based on *ENSDARG00000080337_ACO24175.4* and *tmed1b*. Expression of these markers was significantly higher in the HSPCs II cluster (cl2) than the HSPCs I cluster (cl4) (Figure 5h, S3). Cells in cluster 5 expressed endothelial transcripts, that are known to be involved in the EHT-process (*cdh5*^51,52^*, adgrg1*^43,53^), indicating an EHT-progenitor lineage (Figure 5c, S3). Signature genes *mpx, lyz, marco,* and *mfap4* were expressed in cluster 1 and 3^54^, indicating a myeloid/neutrophil- and myeloid/monocyte-progenitor lineage, respectively (Figure 5d,e, S3). All markers that were used to identify clusters are listed in Table S1 and the distribution of expression of selected markers is depicted in Figure S4.

**Fig. 5.**
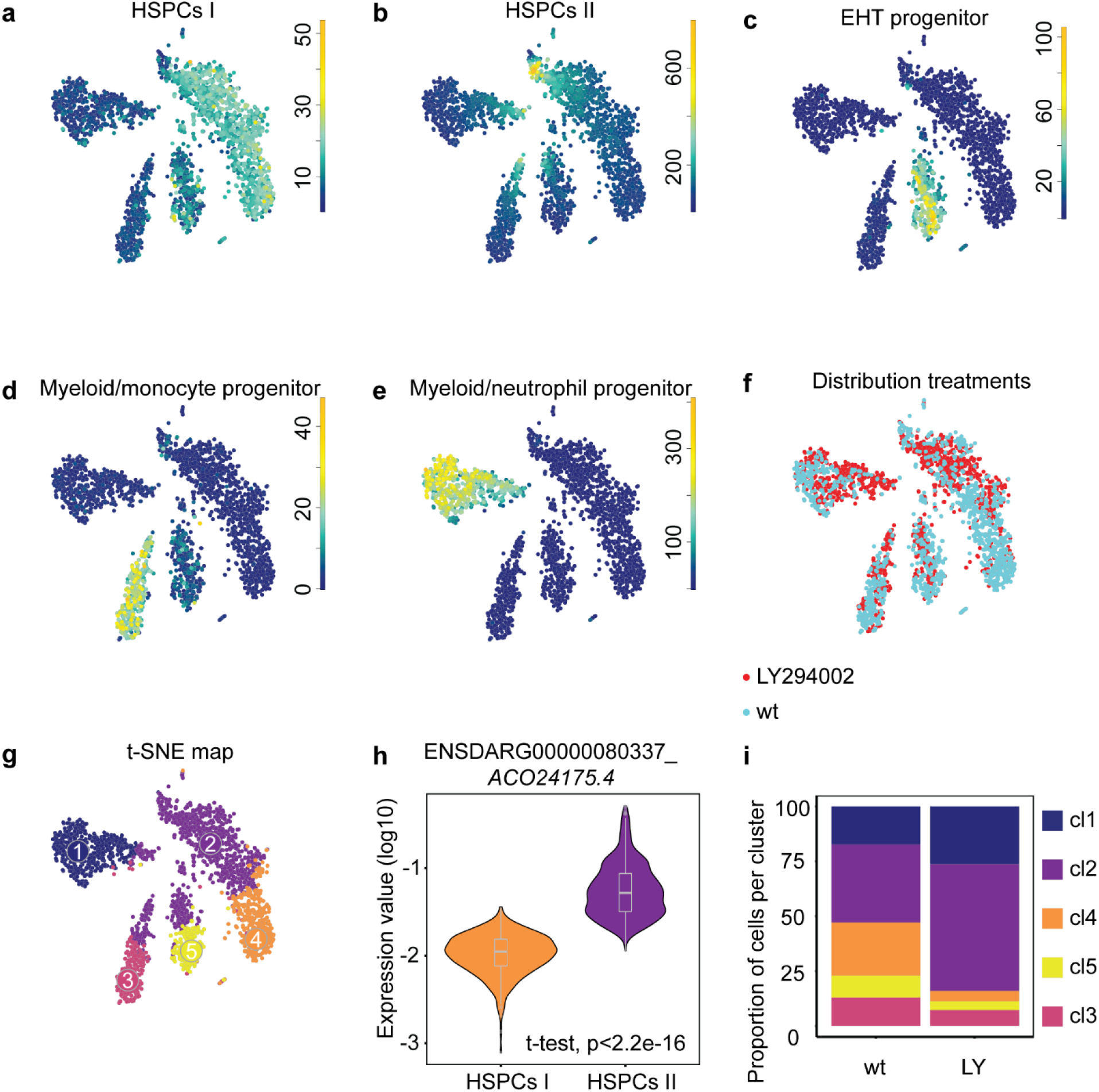
scRNA-seq reveals two types of HSPCs, one of which is lost upon inhibition of the PI3K-pathway. Tissue from control and LY294002-treated embryos (~2,000 each) was dissected, the AGM regions pooled, dissociated and FACS sorted, after which the SORT-seq protocol was performed. (a-e) t-SNE maps highlighting the expression of marker genes for each of the different cell types found. Transcript counts are given in a linear scale. (a) HSPCs I, (b) HSPCs II, (c) EHT progenitor, (d) Myeloid/monocyte progenitors, (e) Myeloid/neutrophil progenitors. (f) t-SNE map highlighting the distribution of cells from LY294002-treated embryos and their controls (g). Visualization of k-medoid clustering and cell-to-cell distances using t-SNEs. Each dot represents a single cell. Colors and numbers indicate cluster and correspond to colors in (i). The distribution of in total 2,512 cells over the five clusters are shown as percentage of total for control and LY294002-treated embryos. Fisher’s Exact test with multiple testing correction (Fdr) were used for statistical analysis. *** p<0.001. cl1: Myeloid/neutrophil progenitor, cl2: HSPC II, cl3: myeloid/monocyte progenitor, cl4: HSPC I, cl5: EHT progenitor.(h) normalized expression level of ENSDARG00000080337_*ACO24175.4* for HSPC I and HSPC II cluster using violin plots. Normalized expression is plotted on a log10 scale.

Cells from LY294002-treated embryos had an uneven distribution over all clusters. In HSPCs II and myeloid/neutrophil progenitors, cells from LY294002-treated embryos were overrepresented compared to control embryos (Fisher’s Exact test, p<0.001). In HSPCs I, EHT- and myeloid/monocyte progenitors, cells from LY294002-treated embryos were underrepresented (Fisher’s Exact test, p<0.001) (Figure 5f,i). These data indicate that LY294002-treatment led predominantly to loss of cells from HSPCs I cluster, which is consistent with the imaging data where half of the HSPCs fail to complete EHT (Figure 3d-g). The loss of HSPCs I in response to PI3K inhibition is accompanied by an increase in HSPCs II and myeloid/neutrophil progenitors.

### More HSPCs upon inhibition of PI3K and less HSPCs in *pten* mutants

The CHTs of approximately 100 control and 100 LY294002-treated embryos were processed for scRNA-seq. Of the 928 cd41^low^ cells that were analyzed, 684 remained after filtering. RaceID3 separated the cells in distinct clusters (Figure 6a). Cells in cluster 2 expressed erythrocyte progenitor-related genes (*hbbe2, alas2* and *cahz*) (Figure 6b). Cluster 3 is characterized by cells expressing genes related to thrombocyte/erythrocyte progenitors (*gata1a, klf1*^50,54,55^) (Figure 6c). Cells in cluster 1 express genes indicative of HSPCs, including *c-myb* (Figure 6d). Cluster 4 represents early myeloid progenitors, as *runx3, pu.1 (*also known as *spi1b)* and *cebpb*^54^ are highly expressed (figure 6e). Cluster 5 is characterized by neutrophil progenitor-related gene expression (*mpx)* (figure 6f). Analysis of the distribution of hematopoietic cells, using a Fisher’s exact test indicated that the thrombocyte/erythrocyte progenitor cells were underrepresented in the LY294002-treated embryos (p<0.01) and HSPCs were significantly overrepresented (p<0.001) (Figure 6g,h).These results indicate a significant shift towards HSPCs at the expense of the thrombocyte/erythrocyte progenitor cluster in response to LY294002 treatment.

**Fig. 6.**
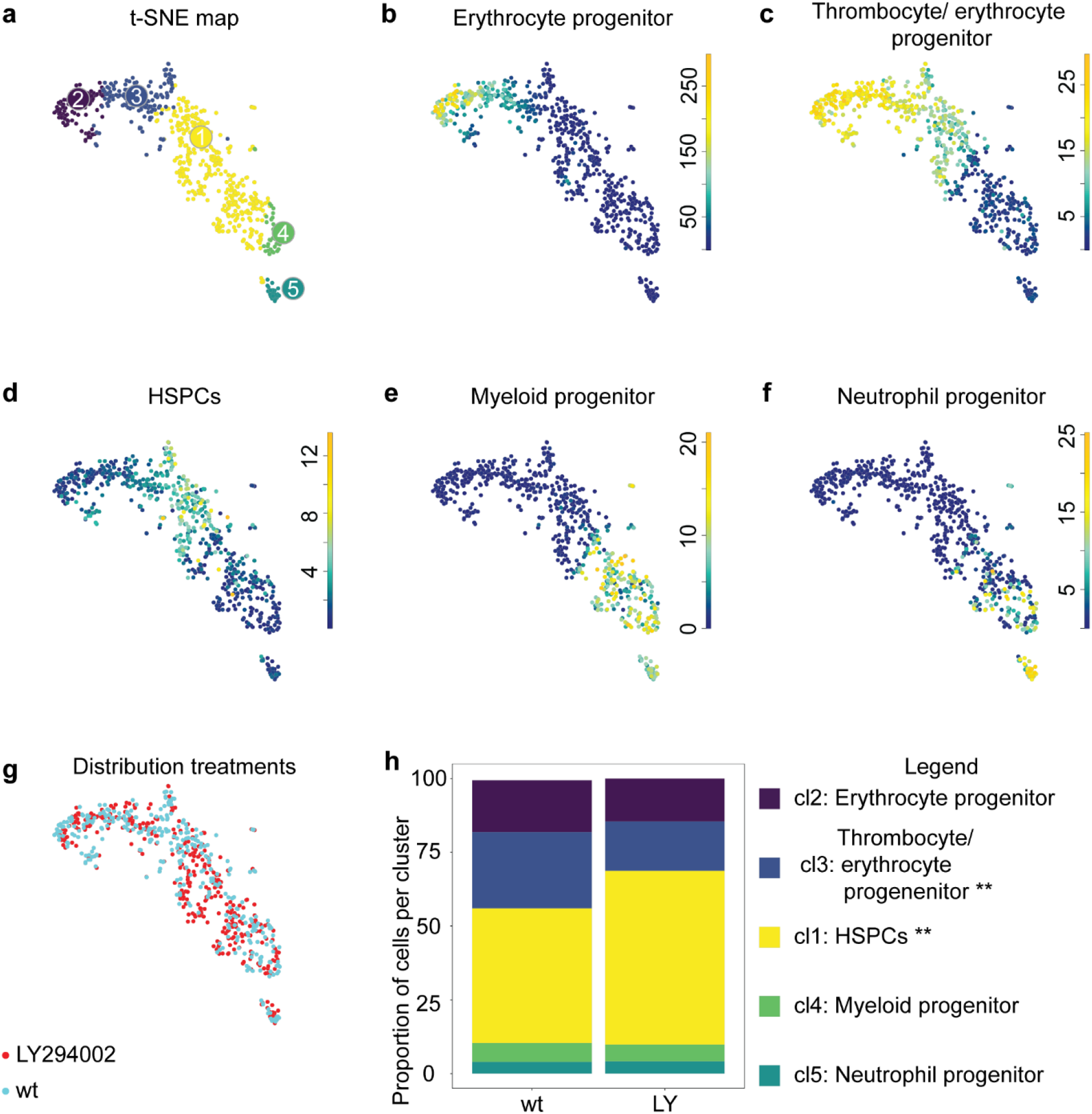
ScRNA-seq seq reveals a shift towards HSPCs in LY294002-treated 5dpf old embryos. CHTs of control and LY294002-treated embryos (5 dpf, ~100 embryos each) were dissected, pooled, dissociated, FACS-sorted and submitted to SORT-seq. (a) Visualization of k-medoid clustering and cell-to-cell distances using t-SNEs. Each dot represents a single cell. Colors and numbers indicate cluster and correspond to colors in (h). In total 684 cells are shown. (b-f) t-SNEs maps highlighting the expression of marker genes for each of the different cell types found. Transcript counts are given in a linear scale. (b) Erythrocyte progenitors, (c), Thrombocyte/erythrocyte progenitors, (d) HSPCs, (e) Myeloid progenitors, (f) Neutrophil progenitors. (g) t-SNE map highlighting the distribution of LY294002-treated embryos and their controls (h). The percentage of cells from LY294002-treated embryos and their controls in the different clusters. Fisher’s Exact test with multiple testing correction (Fdr) were used for statistical analysis. ** p<0.01, *** p<0.001.

Likewise, we assessed transcriptomic differences by scRNA-seq in HSPCs from the CHT between *ptena^−/−^ ptenb^−/−^* mutant embryos and their siblings at 5 dpf. Approximately 100 *ptena^−/−^ptenb^−/−^* mutant embryos and siblings were selected based on phenotype^15^, which yielded 614 cd41^low^ cells after filtering. RaceID3 indicated that clusters emerged representing the same hematopoietic lineages as described for the wild type and LY294002-treated data (cf. Figure 6 and 7). Analysis of the distribution of hematopoietic cells from *pten* mutants and their siblings over the five clusters indicated that the erythrocyte- and neutrophil progenitor cells were overrepresented in the pten mutant (p<0.001 and p<0.05) and that HSPCs were significantly underrepresented (p<0.001)( Figure 7g-h, S5). These results indicate a significant shift in *ptena^−/−^ptenb^−/−^* mutant embryos towards erythrocyte progenitor and neutrophil progenitors at the expense of HSPCs.

**Fig. 7.**
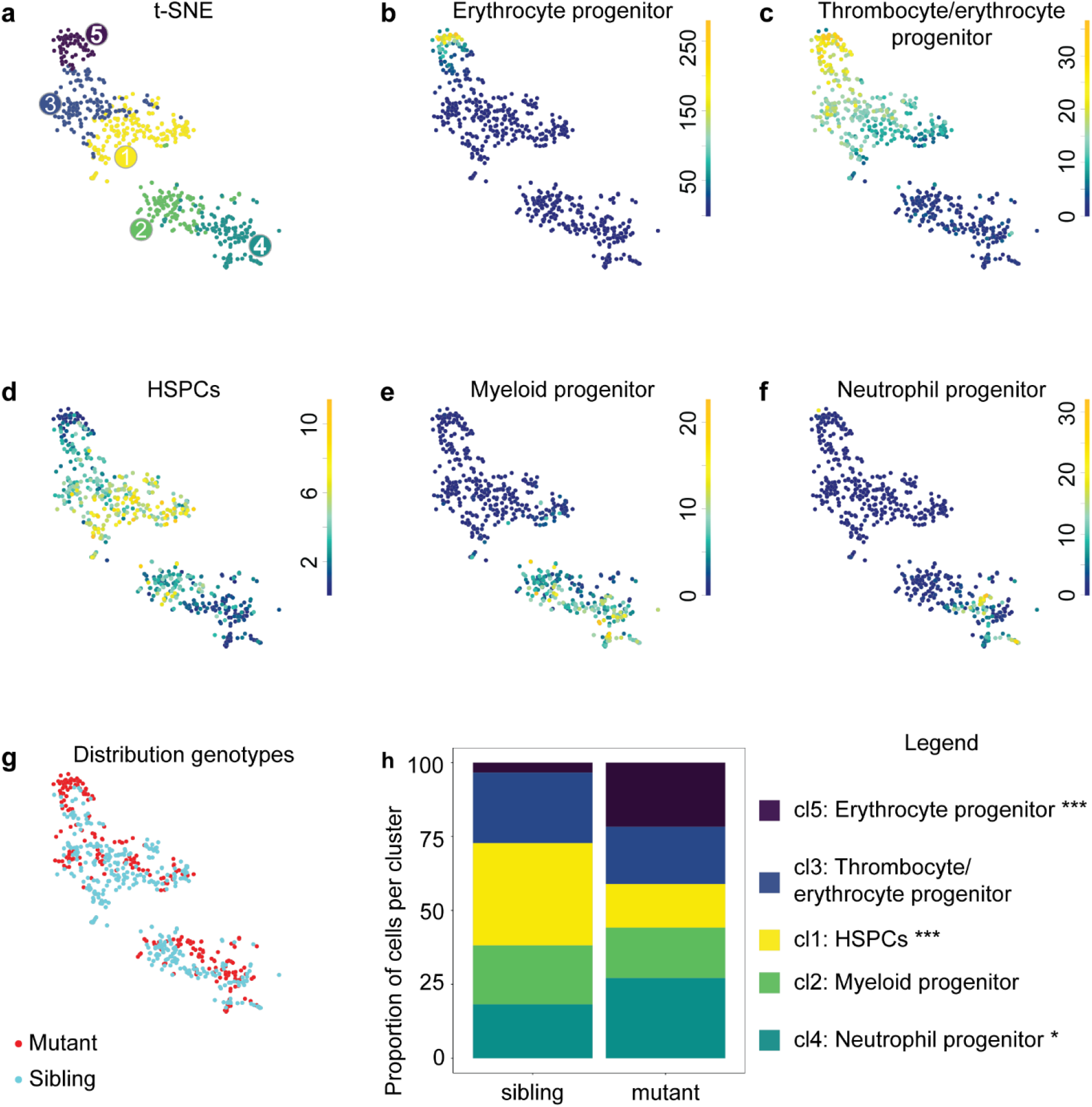
ScRNA-seq seq reveals a shift towards more differentiated cell types in 5 dpf old *ptena^−/−^ptenb^−/−^* mutant embryos. CHTs of control and *ptena^−/−^ptenb^−/−^* mutant embryos (5 dpf, ~100 embryos each) were dissected, pooled, dissociated, FACS-sorted and submitted to SORT-seq. (a) Visualization of k-medoid clustering and cell-to-cells distances using t-SNEs. Each dot represents a single cell. Colors and numbers indicate cluster and correspond to colors in (h). In total 614 cells are shown. (b-f) t-SNEs maps highlighting the expression of marker genes for each of the different cell types found. Transcript counts are given in a linear scale. (b) Erythrocyte progenitors, (c), Thrombocyte/erythrocyte progenitor, (d) HSPCs, (e) Myeloid progenitors, (f) Neutrophil progenitors. (g) t-SNE map highlighting the distribution of *ptena^−/−^ptenb^−/−^* mutant embryos and their siblings. (h). The percentages of cells from *ptena^−/−^ptenb^−/−^* mutant embryos and their siblings in the different clusters. Fisher’s Exact test with multiple testing correction (Fdr) were used for statistical analysis. * p<0.05, *** p<0.001.

## Discussion

We used zebrafish mutant embryos lacking functional Pten to investigate how loss of Pten affects the ontogeny of hematopoiesis. Characterization of zebrafish *ptena^−/−^ptenb^−/−^* mutant embryos led to the unexpected finding that half of the HSPCs undergo apoptosis upon emergence from the VDA during EHT at the onset of the definitive wave (Figure 1). Loss of function of Pten is usually linked to enhanced cell survival, such as for instance in *Pten* knock-out mice^56^. We reported that ɣ-irradiation reduces apoptosis in *ptena^−/−^ptenb^−/−^* mutant embryos^15^. Apoptosis of zebrafish HSPCs has been reported before, in that *grechetto* mutants display decreasing numbers of HSPCs due to apoptosis^51^. Runx1 knockdown also induced abortive EHT events due to apotosis^5^. *Runx1* expression was not affected in the VDA of *ptena^−/−^ ptenb^−/−^* mutants (Figure S1), suggesting that the mechanism underlying EHT defects in *ptena^−/−^ptenb^−/−^* mutant embryos and Runx1 morphants are distinct. Apoptosis of HSPCs in *pten* mutants is due to enhanced PI3K-mediated signaling, because treatment with a PI3K inhibitor rescued apoptosis of HSPCs. Surprisingly, treatment of wild type embryos with the PI3K inhibitor induced death of half of the HSPCs upon emergence from the VDA as well (Figure 3). These results suggest that upon emergence from the VDA, HSPCs require a moderate level of PI3K signaling, as hyperactivation of PI3K signaling in Pten mutants as well as inhibition of PI3K signaling induced apoptosis of emerging HSPCs.

After emerging from the VDA, the surviving HSPCs enter circulation and seed the CHT, as demonstrated by photoconversion of endothelial cells prior to EHT in *ptena^−/−^ptenb^−/−^* mutant embryos and siblings (Figure 2). Half of the HSPCs of *ptena^−/−^ptenb^−/−^* mutant embryos and LY294002-treated embryos colonized the CHT, compared to wild type embryos (Figure 2,3, Table 1). In LY294002-treated embryos the decrease in HSPCs remained, whereas in *ptena^−/−^ptenb^−/−^* mutant embryos the surviving HSPCs hyperproliferate leading to an increase in HSPCs at later stages^17^. Surviving HSPCs from *ptena^−/−^ptenb^−/−^* mutants engage in all blood lineages^17^. However, definitive differentiation of major blood lineages is arrested in the *ptena^−/−^ptenb^−/−^* mutants, consistent with the inverse correlation of proliferation and differentiation of stem cells^57^. The surviving HSPCs of LY294002-treated embryos also engaged in all blood lineages (Figure 4), demonstrating pluripotency of the HSPCs.

Using scRNA-seq at the onset of the definitive wave (36hpf) of hematopoiesis two HSPC clusters were identified, that both expressed HSPC markers. In control embryos equal numbers of cells populated the HSPCs I and HSPCs II clusters. Predominantly the cells from the HSPCs I cluster were lost upon PI3K-inhibition (Figure 5). Our imaging data indicated that half of the HSPCs disintegrated upon treatment with LY294002 (Figure 3). It is tempting to speculate that the surviving half of the HSPCs all belong to the HSPCs II cluster. Whereas both HSPCs clusters expressed HSPC markers, expression of ENSDARG00000080337_*ACO24175.4* and to a lesser extent *tmed1b* distinguished the HSPCs II cluster from the HSPCs I cluster. *In situ* hybridization using an ENSDARG00000080337_*ACO24175.4-*specific probe indicated high expression throughout the embryo, which did not allow validation of the difference in expression in HSPCs I and HSPCs II cells (Figure S3). Little is known about ENSDARG00000080337_*ACO24175.4*, except that it is a mitochondrial ribosomal gene (mt rDNA). Interestingly, HSCs have significantly lower rates of protein synthesis than other hematopoietic cells^58^. The protein product of ENSDARG00000080337_*ACO24175.4* may have a role in protein synthesis. Hence, the difference in expression levels may indicate that the HSPCs II cells that survive PI3K-inhibition are less stem cell-like and more progenitor-like, poised to differentiate.

In response to LY294002-treatment the number of cd41^low^ HSPCs was reduced in the CHT at 4 dpf and in the definitive hematopoietic organs at 8 and 12 dpf (Figure 3,4). scRNA-seq of putative HSPCs (cd41^low^, kdrl^+^ cells) at the end of the definitive wave (5dpf) indicated initiation of differentiation in different blood lineages (Figure 6), consistent with *in situ* hybridization (Figure 4). Yet, inhibition of PI3K arrested differentiation, i.e. increased HSPC fate, predominantly at the expense of thrombocyte/erythrocyte progenitor fate (Figure 6). Overall, it is evident that there is a significant reduction in hematopoietic cell number (Figure 4, 6), which may be caused by preferential loss of HSPCs with more stem cell-like properties (Figure 5).

ScRNA-seq at the end of the definitive wave showed a significant increase in erythrocyte- and neutrophil-progenitors in *ptena^−/−^ptenb^−/−^* mutant embryos (Figure 7, S5), consistent with earlier *in vivo* data^17^. However, we reported an overall increase in HSPCs, due to hyperproliferation, whereas here, we observed a decrease in HSPCs in the scRNA-seq data. An explanation for this apparent discrepancy is that the hyperproliferating HPSCs we observed earlier^17^ actually have initiated differentiation already and are scored as erythrocyte and neutrophil progenitors by scRNA-seq.

Conditional knock-out of *Pten* in HSCs in mouse adult bone marrow, drives HSCs into the cell cycle, resulting in transient expansion of the spleen and eventually in depletion of HSCs in the bone marrow. These conditional PTEN-deficient mice die of a myeloproliferative disorder that resembles acute myeloid/lymphoid leukemia, indicating that PTEN is required for maintenance of HSCs^13,14^. It is noteworthy that there are differences between the conditional mouse models and the zebrafish model we used. In the mouse, Pten is deleted in adult bone marrow cells, well after HSCs have formed, whereas in zebrafish, Pten is systemically deleted and therefore effective prior to the emergence of HSPCs. Studies in mice showed that regardless of cell state, HSCs and multi-potent progenitors had a lower protein synthesis rate than more restricted hematopoietic progenitors. Loss of PTEN in HSCs caused depletion of HSCs, due to a higher rate of protein synthesis^58^, which is consistent with our observation that loss of Pten in zebrafish caused HSPCs to hyperproliferate and become less stem-cell like.

Long-term HSCs are quiescent, whereas short-term HSCs proliferate more^2^. It would be tempting to speculate that the HSPCs that undergo apoptosis upon loss of Pten or upon PI3K-inhibition are involved in long-term colonization of definitive hematopoietic organs. The surviving HSPCs in *pten* mutants at the onset of the definitive wave would then represent multi-potent progenitors that only have limited potential for self-renewal. Investigating the regulatory network underlying the surviving and disintegrating HSPCs will further expand our understanding of short- and long-lived HSCs and will eventually contribute to the development of efficient stem cell based therapies^59,60^.

## Acknowledgements

The authors would like to thank Mark Reijnen and animal caretakers for excellent management of the fish facility. Microscopy was done in the Hubrecht Imaging Centre. The authors would like to thank Stefan van der Elst and Reinier van der Linden for FACS-sorting. The authors would like to thank Jeroen Paardekooper Overman for statistical analysis, Laila Ritsma, Sylvain de Rossi and Miriam Stumpf for technical support and Bas Castelijns for help with scRNA data analysis. This work was supported in part by an EU (FP7) grant, ZF-CANCER (HEALTH-F2-2008-201439).

## Authorship

S.B.F, S.C. and J.d.H. designed experiments with input from K.K. and P.H.; S.B.F and S.C. performed the experiments; B.P. and S.S.M. generated the Tg(kdrl:Dendra) line and helped perform the photoconversion experiments; K.K. and J.d.H. supervised the work; S.B.F, S.C., P.H., K.K. and J.d.H. wrote the manuscript.

Correspondence and requests for materials should be addressed to: Jeroen den Hertog, Hubrecht Institute, Uppsalalaan 8, 3584 CT Utrecht, the Netherlands.

