## Supplementary material for "Phosphatidylinositol-3 kinase activity controls survival and stemness of zebrafish hematopoietic stem/progenitor cells": Suppl Table S1 and Figures S1-S5

Table S1. Marker genes used for identification of clusters in single cell RNA seq. ENSDARG numbers and gene names are given.

| Cluster/Cell type | Gene | ENSDARG |
|---|---|---|
| Thrombocyte/erythrocyte progenitors | *hmbsb* | ENSDARG00000055991 |
| | *prdx3* | ENSDARG00000032102 |
| | *urod* | ENSDARG00000006818 |
| | *uros* | ENSDARG00000027491 |
| | *tubb1* | ENSDARG00000053066 |
| | *klf1* | ENSDARG00000017400 |
| | *gata1a* | ENSDARG00000013477 |
| | *epb41b* | ENSDARG00000029019 |
| Erythrocyte progenitors | *alas2* | ENSDARG00000038643 |
| | *cahz* | ENSDARG00000011166 |
| | *hbbe2* | ENSDARG00000045143 |
| | *hbae3* | ENSDARG00000079305 |
| | *rhag* | ENSDARG00000019253 |
| | *epor* | ENSDARG00000090834 |
| HSPCs | *myb* | ENSDARG00000053666 |
| | *her6* | ENSDARG00000006514 |
| | *ahcy* | ENSDARG00000005191 |
| | *pmp22b* | ENSDARG00000060457 |
| | *ncl* | ENSDARG00000002710 |
| | *adh5* | ENSDARG00000080010 |
| | *hmga1a* | ENSDARG00000028335 |
| | *fbl* | ENSDARG00000053912 |
| | *mycb* | ENSDARG00000007241 |
| | *dkc1* | ENSDARG00000016484 |
| | *pes* | ENSDARG00000018902 |
| | *meis1b* | ENSDARG00000012078 |
| | *tal1 (scl)*[49] | ENSDARG00000019930 |
| | *gata2b* | ENSDARG00000009094 |
| | *gfi1aa* | ENSDARG00000020746 |
| | *adgrg1* | ENSDARG00000027222_ |

| | | |
|---|---|---|
| Myeloid progenitor | *crema* | ENSDARG00000023217 |
| | *itm2bb* | ENSDARG00000041505 |
| | *runx3* | ENSDARG00000052826 |
| | *cebpb* | ENSDARG00000042725 |
| | *cxcr4b* | ENSDARG00000041959 |
| | *zfp36l1a* | ENSDARG00000016154 |
| | *coro1a* | ENSDARG00000054610 |
| | *nr4a3* | ENSDARG00000055854 |
| | *pu.1 (spi1b)* | ENSDARG00000000767 |
| Neutrophil progenitor | *cpa5* | ENSDARG00000021339 |
| | *lect2l* | ENSDARG00000033227 |
| | *npsn* | ENSDARG00000010423 |
| | *sms* | ENSDARG00000008155 |
| | *abcb9* | ENSDARG00000056200 |
| | *ch25hl2* | ENSDARG00000038728 |
| | *mpx* | ENSDARG00000019521 |
| | *lyz* | ENSDARG00000057789 |
| | *srgn* | ENSDARG00000077069 |
| | *mmp9* | ENSDARG00000042816 |
| Monocyte progenitor | *marco* | ENSDARG00000059294 |
| | *ctss2.2* | ENSDARG00000013771 |
| | *mfap4* | ENSDARG00000090783 |
| | *marckls1a* | ENSDARG00000039034 |
| | *ctsba* | ENSDARG00000055120 |
| | *cxcr3.3* | ENSDARG00000070669 |
| | *timp2b* | ENSDARG00000075261 |
| | *lgmn* | ENSDARG00000039150 |
| | *ndrg1a* | ENSDARG00000032849 |
| EHT markers | *edn2* | ENSDARG00000017255 |
| | *efna1b* | ENSDARG00000018787 |
| | *cdh5* | ENSDARG00000075549 |
| | *krt18* | ENSDARG00000018404 |
| | *krt8* | ENSDARG00000058358 |
| | *dab2* | ENSDARG00000031761 |
| | *serpinh1b* | ENSDARG00000019949 |
| | *anxa2a* | ENSDARG00000003216 |
| | *ctsla* | ENSDARG00000007836 |
| | *hapln1b* | ENSDARG00000068516 |
| | *clic2* | ENSDARG00000010625 |
| | *cd81a* | ENSDARG00000036080 |
| | *tie1* | ENSDARG00000004105 |
| HSPCs II 36hpf | *AC024175.4* | ENSDARG00000080337 |
| | *tmed1b* | ENSDARG00000017255 |

### Supplemental Figures

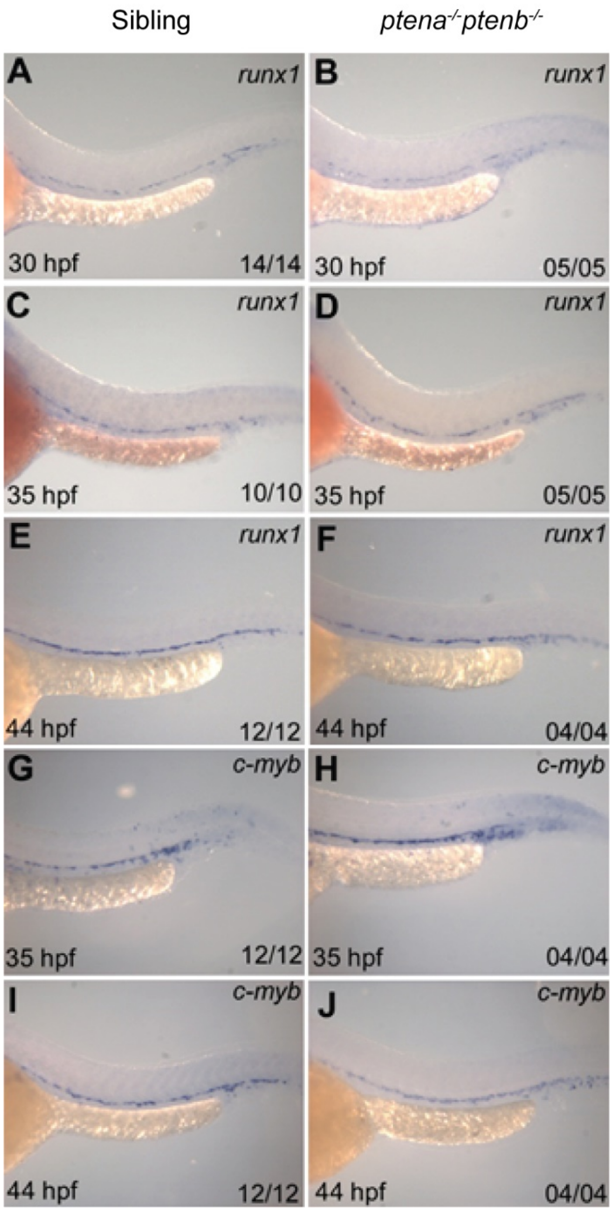


Figure S1. Hemogenic endothelium markers are present in ptena-/-ptenb-/- mutants during onset of definitive hematopoiesis. *Ptena*$^{+/-}$*ptenb*$^{-/-}$ fish were incrossed and embryos were fixed at different time points as indicated (30,35 and 44 hpf). *In situ* hybridization using HSPC markers *runx1* (a-f) and *c-myb* (g-j) was done, pictures were taken and subsequently the genotypes of these embryos was established by sequencing. No differences were observed between *ptena*$^{-/-}$*ptenb*$^{-/-}$ mutant embryos and siblings. Representative embryos are depicted with anterior the left; the number of embryos that showed a particular pattern/total number of embryos is indicated in the bottom right corner of each panel.

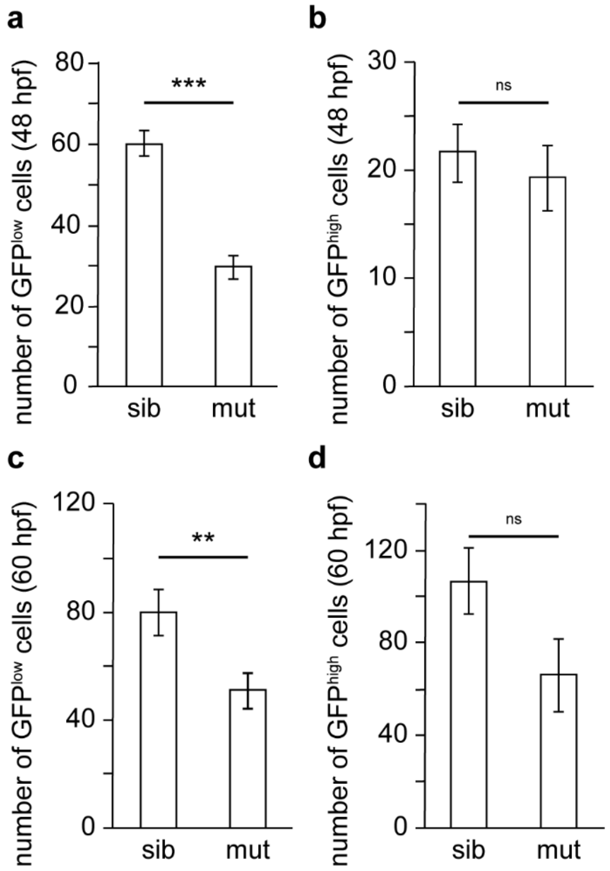


Figure S2. Reduced number of GFP$^{low}$, but not GFP$^{high}$ cells in the CHT of tg(*cd41:eGFP) ptena$^{-/-}$ptenb$^{-/-}$* mutant embryos, compared to siblings. (a) the number of GFP$^{low}$ HSPCs and (b) GFP$^{high}$ thrombocytes at 48 hpf in the CHT of tg(*cd41:eGFP*) siblings (sib) and *ptena$^{-/-}$ptenb$^{-/-}$* mutants (mut), expressed as average number of cells. Error bars indicate standard error of the mean (SEM). (c,d) as in (a,b), but at 60 hpf. Note that the difference in GFP$^{low}$ HSPCs is smaller due to enhanced proliferation and the apparent difference in GFP$^{high}$ cells is almost significant, due to an arrest in differentiation. Shapiro Wilk Test for normal distribution and two-tailed t-test were used for statistical analysis; p-values are: (a) $2.24*10^{-8}$ (***), (b) 0.57 (not significant, ns), (c) 0.013 (**), (d) 0.066 (not significant, ns).

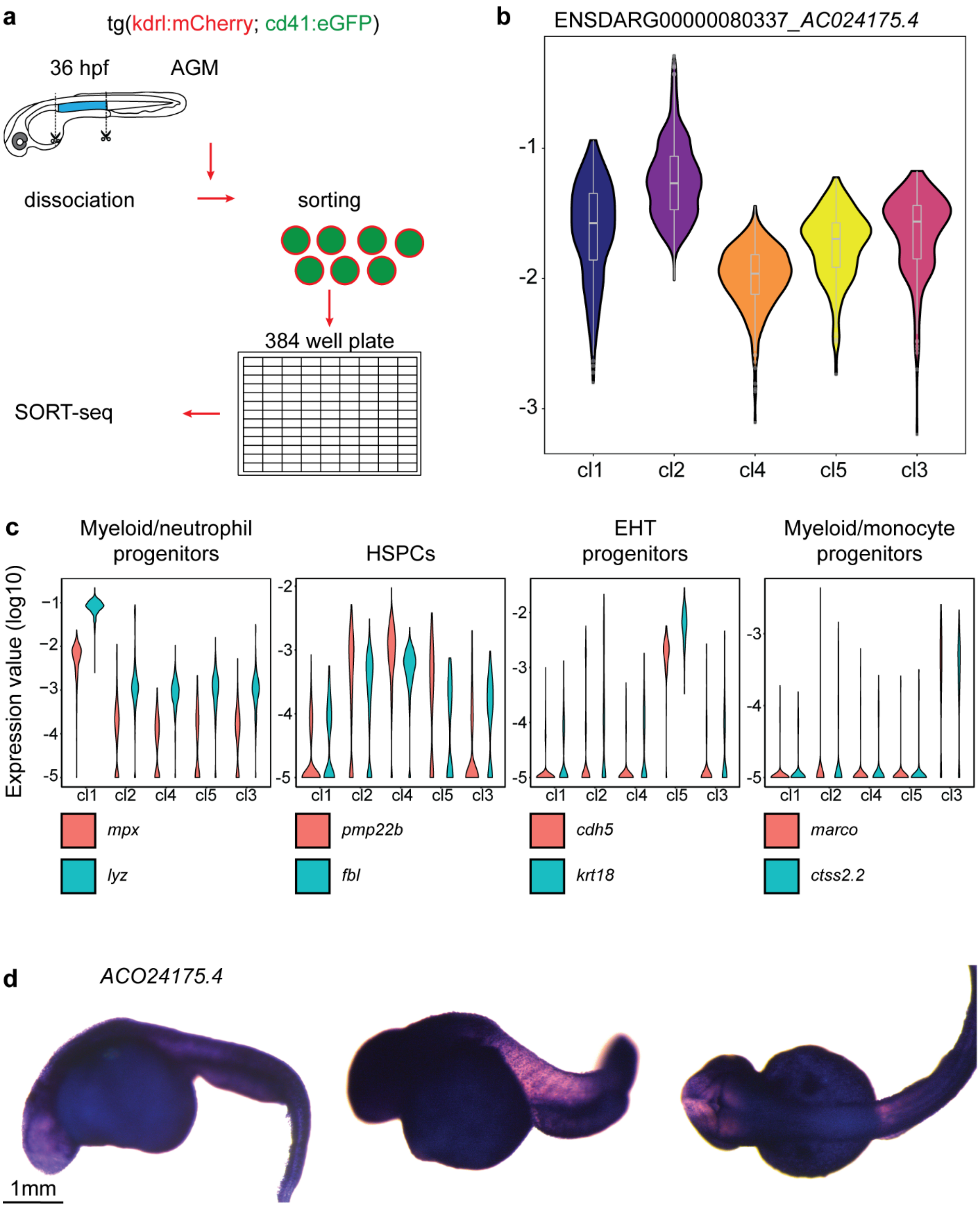
a
tg(kdrl:mCherry; cd41:eGFP)
36 hpf
AGM
dissociation
sorting
384 well plate
SORT-seq
b
ENSDARG00000080337_AC024175.4
-1
-2
-3
cl1
cl2
cl4
cl5
cl3
c
Myeloid/neutrophil progenitors
HSPCs
EHT progenitors
Myeloid/monocyte progenitors
Expression value (log10)
mpx
lyz
pmp22b
fbl
cdh5
krt18
marco
ctss2.2
d
ACO24175.4
1mm

Figure S3. Single cell RNA seq of control and PI3K inhibitor treated embryos at the onset of definitive hematopoiesis. (a) Workflow of scRNA seq. Tissue from control and LY294002-treated embryos (~2,000 each) was dissected, the AGM regions pooled, dissociated and FACS sorted, after which the SORT-seq protocol was performed. (b) Normalized expression of ENSDARG00000080337_*ACO24175.4* and *tmed1b* over all clusters. Normalized expression is plotted on log10 scale using violin plots and boxplots. cl1: Myeloid/neutrophil progenitor, cl2: HSPC II, cl4: HSPC I, cl5: EHT progenitor, cl3: myeloid/monocyte progenitor. (c) Normalized expression of signature genes for cluster identities using violin plots. Normalized expression value is plotted on a log10 scale. (d) whole mount ISH of 36 hpf wild type embryos using a probe specific for ENSDARG00000080337_*ACO24175.4*. Forward primer: 5'TTAAAGCCCCGAATCCAGGT 3', reverse primer with T7 promoter: GAGTAATACGACTCACTATAGGTTTTGGTAAACAGGCGAGGC. At this stage, this gene is expressed throughout the embryo at a very high level, which does not allow to distinguish between individual blood cells.

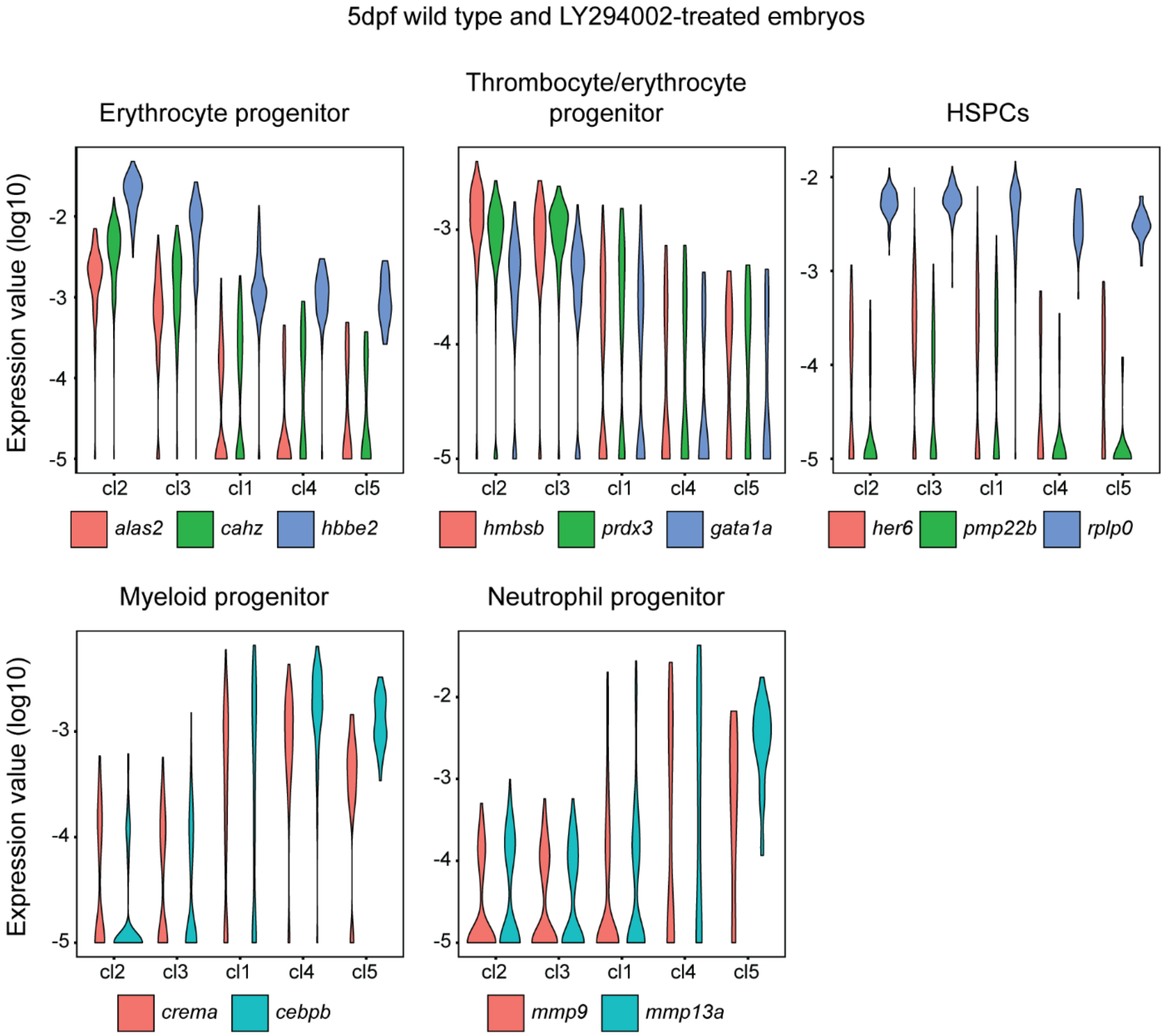


Figure S4. Cluster identities at 5 dpf for wild type and LY294002-treated embryos. Normalized expression of signature genes for cluster identities using violin plots. Normalized expression value is plotted on a log10 scale.

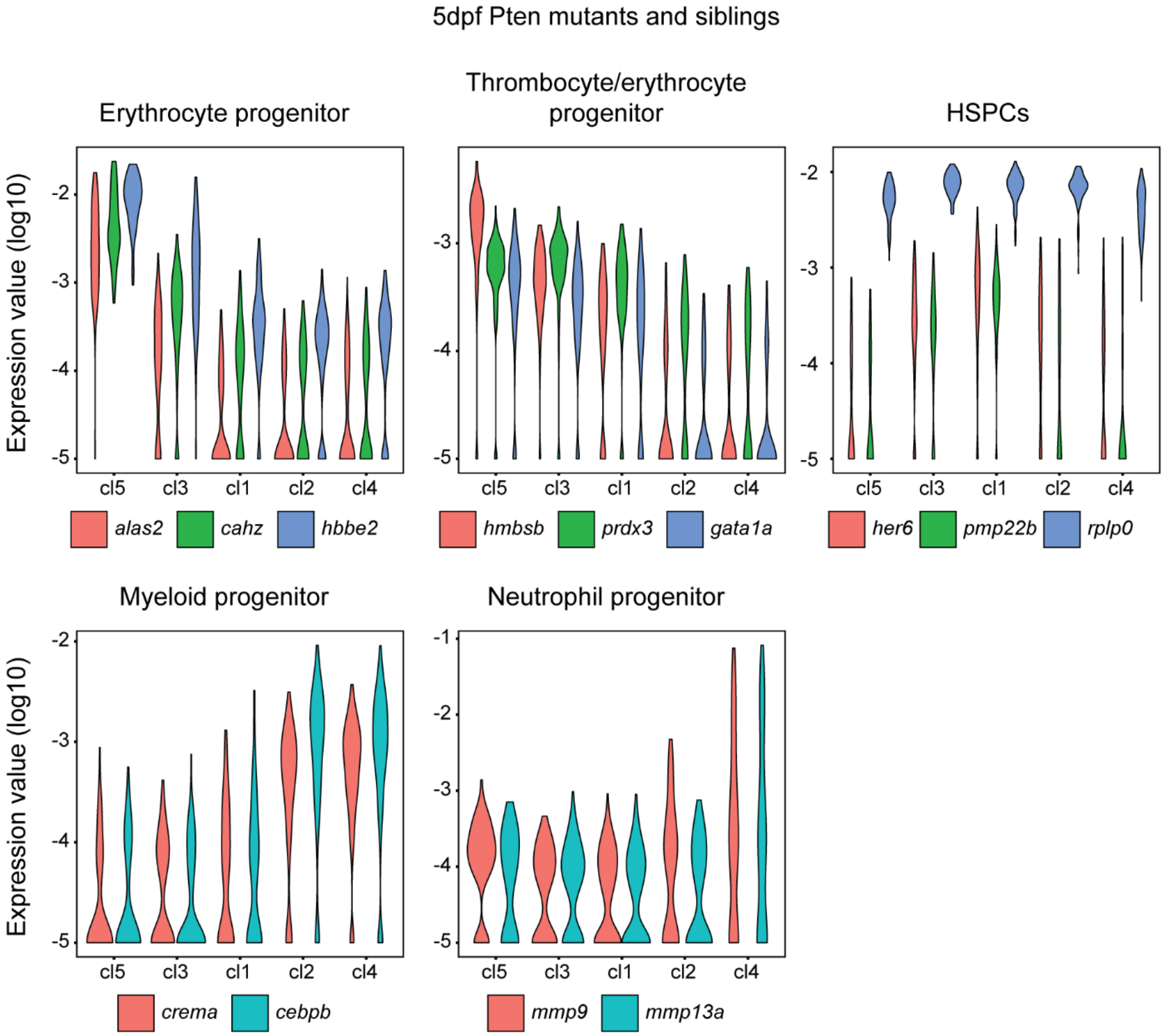


Figure S5. Cluster identities at 5 dpf for ptena-/-ptenb-/- mutant embryos and their siblings. Normalized expression of signature genes for cluster identities using violin plots. Normalized expression value is plotted on a log10 scale.

#### Supplemental Movies

Movie S1. Time lapse confocal imaging of endothelial to hematopoietic transition in ptena$^{-/-}$ptenb$^{-/-}$ zebrafish mutant embryo in *tg(kdrl-gfp)* transgenic background, imaged from 35 hpf onwards. Cell tracks of cells undergoing endothelial to hematopoietic transition are indicated in color. Time is indicated in hours.

Movie S2. Time lapse confocal imaging of endothelial to hematopoietic transition in *tg(kdrl-gfp)* transgenic background zebrafish embryo treated with LY294002 (5 µM) from 32 hpf onwards. Imaging was done from 35 hpf onwards. Cell tracks of cells undergoing endothelial to hematopoietic transition are indicated in color. Time is indicated in hours.
